# Building a schizophrenia genetic network: Transcription Factor 4 regulates genes involved in neuronal development and schizophrenia risk

**DOI:** 10.1101/215715

**Authors:** Hanzhang Xia, Fay M. Jahr, Nak-Kyeong Kim, Linying Xie, Andrey A. Shabalin, Julien Bryois, Douglas H. Sweet, Mohamad M. Kronfol, Preetha Palasuberniam, MaryPeace McRae, Brien P. Riley, Patrick F. Sullivan, Edwin J. van den Oord, Joseph L. McClay

**Affiliations:** Center for Biomarker Research and Precision Medicine, Virginia Commonwealth University, Richmond, Virginia, United States.; Department of Pharmacotherapy and Outcomes Science, Virginia Commonwealth University, Richmond, Virginia, United States.; Department of Biostatistics, Virginia Commonwealth University, Richmond, Virginia, United States.; Department of Psychiatry, University of Utah, Salt Lake City, Utah, United States.; Department of Medical Epidemiology and Biostatistics, Karolinska Institute, Stockholm, Sweden.; Department of Pharmaceutics, Virginia Commonwealth University, Richmond, Virginia, United States.; Virginia Institute for Psychiatric and Behavioral Genetics, Virginia Commonwealth University, Richmond, Virginia, United States.; Departments of Genetics and Psychiatry, University of North Carolina School of Medicine, Chapel Hill, North Carolina, United States.

## Abstract

The transcription factor 4 (*TCF4*) locus is a robust association finding with schizophrenia (SZ), but little is known about the genes regulated by the encoded transcription factor. Therefore, we conducted chromatin immunoprecipitation sequencing (ChIP-seq) of TCF4 in neural-derived (SH-SY5Y) cells to identify genome-wide TCF4 binding sites, followed by data integration with SZ association findings. We identified 11,322 TCF4 binding sites overlapping in two ChIP-seq experiments. These sites are significantly enriched for the TCF4 Ebox binding motif (>85% having ≥1 Ebox) and implicate a gene set enriched for genes down-regulated in TCF4 siRNA knockdown experiments, indicating the validity of our findings. The TCF4 gene set was also enriched among 1) Gene Ontology categories such as axon/neuronal development, 2) genes preferentially expressed in brain, in particular pyramidal neurons of the somatosensory cortex, and 3) genes down-regulated in post-mortem brain tissue from SZ patients (OR=2.8, permutation p<4x10^−5^). Considering genomic alignments, TCF4 binding sites significantly overlapped those for neural DNA binding proteins such as FOXP2 and the SZ-associated EP300. TCF4 binding sites were modestly enriched among SZ risk loci from the Psychiatric Genomic Consortium (OR=1.56, p=0.03). In total, 130 TCF4 binding sites occurred in 39 of the 108 regions published in 2014. Thirteen genes within the 108 loci had both a TCF4 binding site ±10kb and were differentially expressed in siRNA knockdown experiments of TCF4, suggesting direct TCF4 regulation. These findings confirm TCF4 as an important regulator of neural genes and point towards functional interactions with potential relevance for SZ.

## INTRODUCTION

Large-scale genome-wide assocation studies (GWAS) have converged on specific risk loci for schizophrenia (SZ) (1). One of the most robust findings is the *TCF4* (transcription factor 4) region on chromosome 18q21.2 (2). First discovered in GWAS meta-analysis (3), the finding remained significant in a follow-up study(4) and a large family-based replication study (5). Most pertinently, SNPs at *TCF4* were among the top findings (p=3.34 x 10^−12^) in the 2014 Psychiatric Genomics Consortium (PGC) mega-analysis of SZ (1). Congruent with the association with SZ, *TCF4* has also been associated with SZ endophenotypes such as neurocognition and sensorimotor gating (6–8).

The biology of *TCF4* suggests a plausible role in CNS disorders: 1) *TCF4* encodes a transcription factor abundantly expressed in brain that has been implicated in neuronal development (9) and function (10, 11); 2) Mutations at *TCF4* cause Pitt-Hopkins Syndrome (PHS), a rare genetic disorder characterized by neurological deficits including mental retardation (2, 12–14); 3) Balanced chromosomal rearrangements in patients with neurodevelopmental disorders have encompassed *TCF4* (15); 4) *TCF4* is a target for transcriptional regulation by microRNA 137 (gene ID: *MIR137)* (16), which is also a top association finding for SZ (1). 5) Transgenic mice that overexpress *TCF4* have cognitive and sensorimotor impairments (17), which mirror deficits observed in SZ patients. Overall, the biological rationale linking *TCF4* to the CNS and SZ is compelling (2), suggesting that further study of this gene could advance our understanding of SZ pathogenesis.

The protein encoded by *TCF4* is a basic helix-loop-helix (bHLH) transcription factor (TF) that recognizes an Ephrussi-box ('E-box') binding site ('CANNTG') (2, 9, 18). However, this motif is too small and non-specific to accurately predict *TCF4* binding computationally. A precise map of binding sites is vital for deciphering the gene regulatory networks under the influence of a TF (19). In recent years, systematic mapping of TF binding has been enabled via chromatin immunoprecipitation coupled to next-generation sequencing (ChIP-seq) (20). ChIP-seq works by precipitating the desired protein-DNA complex out of cell lysate using an antibody complementary to the protein of interest. After removal of the protein, the liberated DNA fragments are sequenced and mapped back to the reference genome to yield a map of regions bound by the protein (21) (Figure 1a).

**Figure 1.**
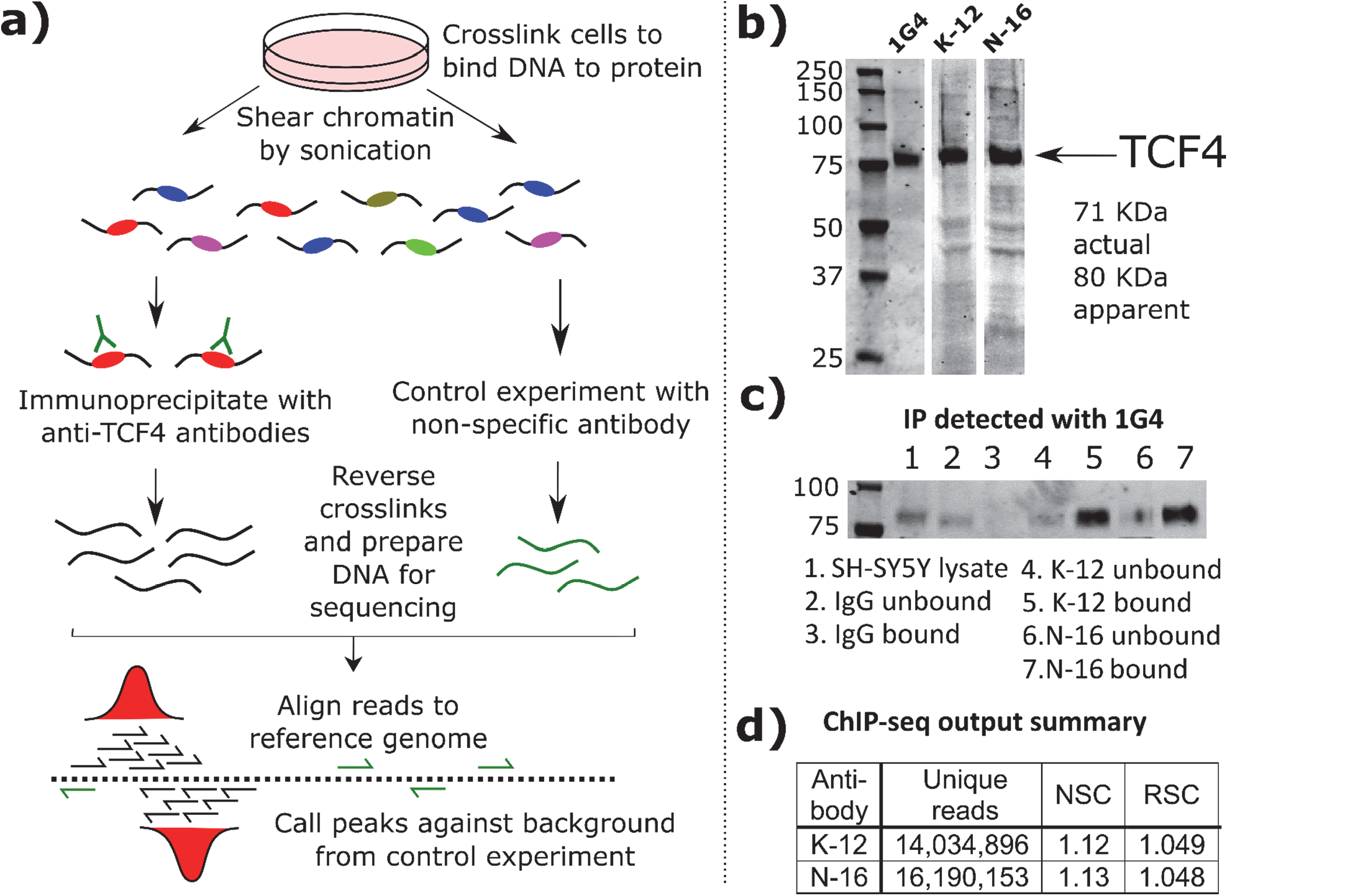
ChIP-seq outline and quality control a. Flow diagram showing the ChIP-seq procedure in cultured cells.
b. Western blotting in SH-SY5Y cell extract using three different anti-TCF4 antibodies (1G4, K-12 and N-16) identifies a single clear band for TCF4 (full length isoform B, UniProt P15884, 667 aa).
c. Crossover IP for TCF4. In this experiment, anti-TCF4 antibodies K-12 and N-16, plus control IgG, were used to immunoprecipitate proteins that were subsequently probed in a Western blot using anti–TCF4 antibody 1G4. Using different anti-TCF4 antibodies in the IP and Western blotting increases confidence that TCF4, rather than a non-specific protein, is detected. The presence of the TCF4 band in the bound (IP) fraction, and not in the bound IgG (control) fraction, indicates that the K-12 and N-16 antibodies successfully immunoprecipitated TCF4.
d. ChIP-seq output summary. The number of uniquely aligning reads for each ChIP-seq experiment are shown. The Normalized Strand Cross-correlation coefficient (NSC) and Relative Strand Crosscorrelation coefficient (RSC) values indicate enrichment in ChIP (Figure 1d), with higher values indicating more enrichment. NSC values less than 1.1 are relatively low and the minimum possible value is 1 (no enrichment). The minimum possible RSC value is 0 (no signal), highly enriched experiments have values greater than 1, and values much less than 1 may indicate low quality (see genome.ucsc.edu/ENCODE/qualityMetrics.html). The values for our experiments indicate good enrichment.

The ENCODE Consortium, which aims to map all regulatory elements in the human genome, has used ChIP-seq extensively to map binding profiles of many TFs in human cell lines (22). TF binding is tissue-specific, so separate experiments are required to characterize binding in cells from each tissue of interest. At its outset, ENCODE did not have a major CNS focus (23). The consortium attempted to map TCF4 binding sites in bone marrow-derived K562 cells, (https://www.encodeproject.org/experiments/ENCSR000FCF/). However, these data were revoked shortly after release. Here we describe TCF4 ChIP-seq in a CNS-derived cell line. To probe the relationship with SZ, we tested the TCF4 gene network for overlap with SZ risk genes from GWAS and gene expression studies.

### A note on nomenclature

Transcription Factor 4, located on chromosome 18, was previously known by the aliases *E2-2, ITF-2, PTHS*, and *SEF-2* (http://www.ncbi.nlm.nih.gov/gene/6925). *TCF4* and *TCF7L2* (Gene ID: 6934) are often confused because they share the *TCF4* alias (2, 18). *TCF7L2*, located on chromosome 10, encodes “T-Cell Factor 4”, an effector of Wnt/β-catenin signaling (24), that is not an accepted risk factor for SZ. In this study, all references to *TCF4* are to the gene with coordinates chr18:52889562-53332018 (hg19) and these identifiers: HUGO name=TCF4, ENTREZ=6925, HGNC=11634, ENSEMBL=ENSG00000196628, and UNIPROT=P15884.

## RESULTS

### ChIP-seq and peak calling

At the start of the project, no validated ChIP antibodies were available for TCF4. Of eight candidate anti-TCF4 antibodies identified, three passed initial immunoblot testing according to ENCODE guidelines, whereby a single band of the appropriate mass (TCF4-B long isoform, 667 aa) accounting for >50% of the total lane intensity was observed (25) (Figure 1b). We performed IP cross-reactivity studies using these antibodies, where IP with anti-TCF4 antibodies was used as the substrate for Western blot with a different anti-TCF4 antibody. Figure 1c shows that anti-TCF4 antibodies K-12 and N-16 immunoprecipitated a protein of the appropriate mass that was detected by monoclonal anti-TCF4 antibody 1G4, indicating that all three antibodies were likely detecting the same protein. We also conducted mass spectrometry-based proteomics analysis of the IP of all three antibodies and successfully detected TCF4 peptides in each case (Supplementary Figures S1 and S2), while we did not detect any in IgG control IP experiments. We therefore proceeded with ChIP-seq using these antibodies.

Each ChIP experiment involved isolating TCF4-bound DNA from approximately 1.2x10^7^ SH-SY5Y cells. Average DNA yield was within expected parameters (∼10 ng per replicate), but DNA was also recovered from ChIP using the control IgG antibodies. This indicated that non-specific material was captured and that ChIP-seq using the IgG control antibody was the appropriate background to call peaks. After sequencing and alignment to hg19, potential PCR duplicate reads were aggressively filtered (‘rmdup’ command in Samtools), to collapse all reads with the same start position to single reads. Only two of the three antibodies (K-12 and N-16) worked well. Figure 1d shows the summary statistics for these assays and cross-correlation plots are provided in Supplementary Figures S3 and S4. After filtering out ENCODE blacklisted regions, we examined overlap between peaks called for antibodies K-12 and N-16. First we plotted the difference in starting position between peaks observed in K-12 ChIP-seq and the closest peak in N-16 ChIP-seq. Supplementary Figure S5 shows that there was a clear enrichment for peaks with start sites ± 200 bp in each experiment. This is the approximate peak size (∼225 bp) in the ‘narrowpeak’ output format used by the SPP peak caller. Therefore, we specified that the peaks called in the two separate experiments had to overlap (at least 1bp in common) to count as “replicated”; neighboring, non-overlapping peaks were not considered. Applying this criterion resulted in 11,322 TCF4 binding sites present in both experiments, using false discovery rate (FDR) threshold < 1%. As expected, overlap was greater for peaks called with higher confidence. Sorting on SPP signalValue, 65% of peaks in the top 100 in the K-12 experiment had a matching peak in the N-16 data; 50% in the top 250; 45% in the top 500; and 29% in the top 5000.

Supplementary Figure S6 shows the distribution of all 11,322 binding sites by chromosome. No binding sites were obtained for the chrY because SH-SY5Y is genetically female. The genomic distribution was nonrandom with relatively large numbers of peaks detected on chrs 7 and 17. All 11,322 binding sites are provided in Supplementary Table S1.

As a further validation step, we tested for enrichment of specific sequence motifs at the TCF4 binding sites. TCF4 binds the ‘Ebox’ sequence motif (‘CANNTG’). Figure 2 shows the most significantly enriched sequence motifs at the consensus 11,322 ChIP-seq peaks. The top overall motif was the binding site for STAT1 (‘RGRAA‘), while the top ranked TCF4 Ebox (‘CAYCTG’) was fourth. The redundancy of the central 2 bp of the Ebox motif means that several different sequence combinations are possible and, for example, the ‘CACCTG’ Ebox variant was detected at E= 2.3e-016. Overall, 86.4% of the 11,322 binding sites encompassed an Ebox motif.

**Figure 2.**
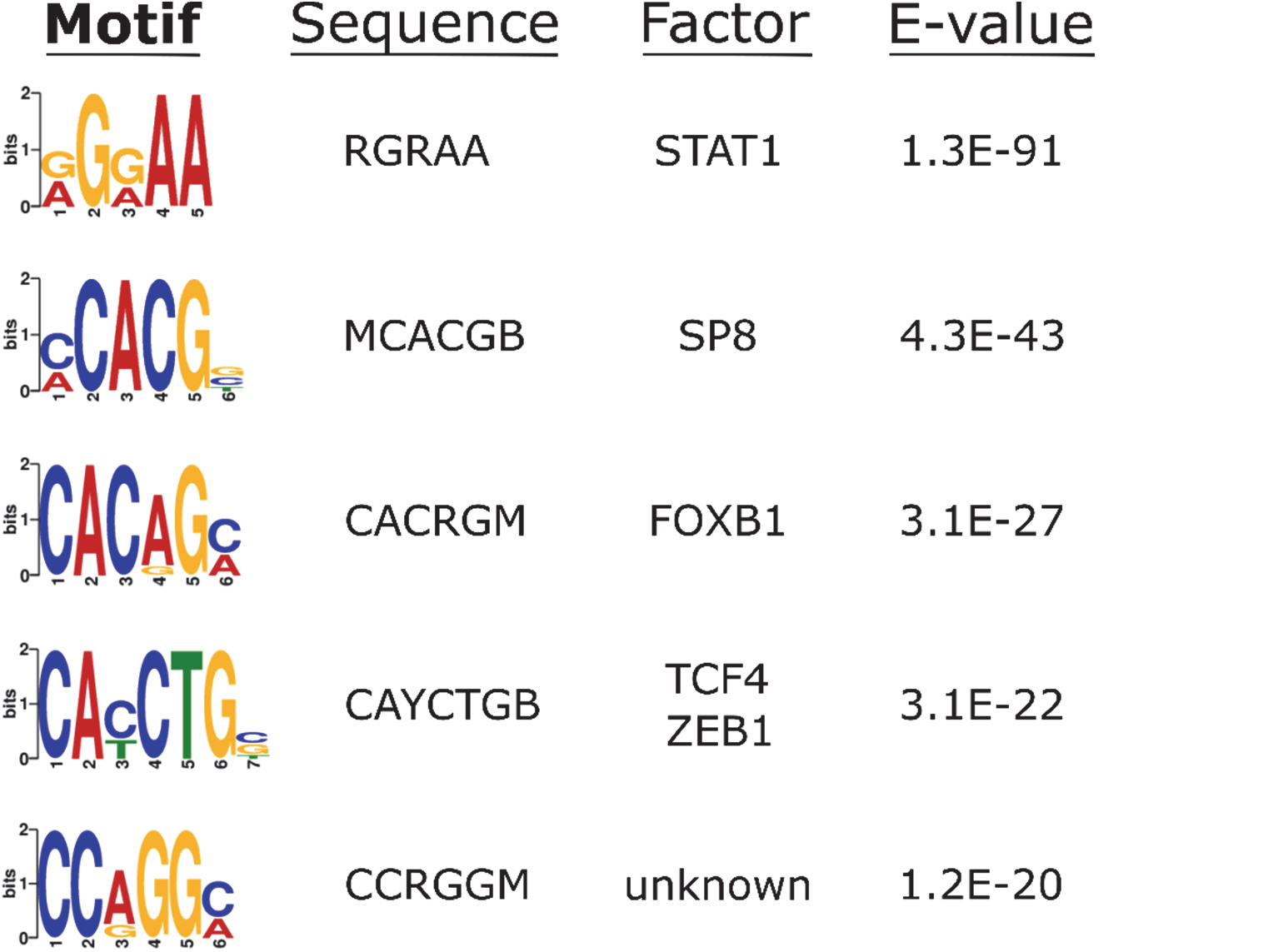
Sequence motif enrichment at TCF4 binding sites. Output from DREME shows the most enriched motifs around the TCF4 ChIP peaks. Motif sequences are shown with conventional nucleotide redundancy codes. E-values are output from DREME/MEME-ChIP and indicate the p-value multiplied by the number of instances tested as a correction for multiple testing.

### Genes associated with TCF4 binding sites and their tissue-specific expression

8923 of the 11322 binding sites were within 10 kb of a RefSeq gene. To obtain insight into TCF4’s functional role, we used GREAT (26) to conduct enrichment tests for GO categories and MSigDB pathways. GREAT first assigns genomic loci to regulatory domains associated with genes and only 34 TCF4 binding sites could not be assigned to any gene (Supplementary Figure S6). The fact that over 2,000 binding sites were not within 10 kb of a gene, yet only 34 could not be assigned to a gene suggests that a number of binding sites are at enhancers or long-range *cis*-regulatory elements. The results from GREAT are shown in Table 1. Significant results in the GO cellular component category had a strong neuronal theme, while insulin signaling and axon/neuronal development were notable pathway findings.

**Table 1.**
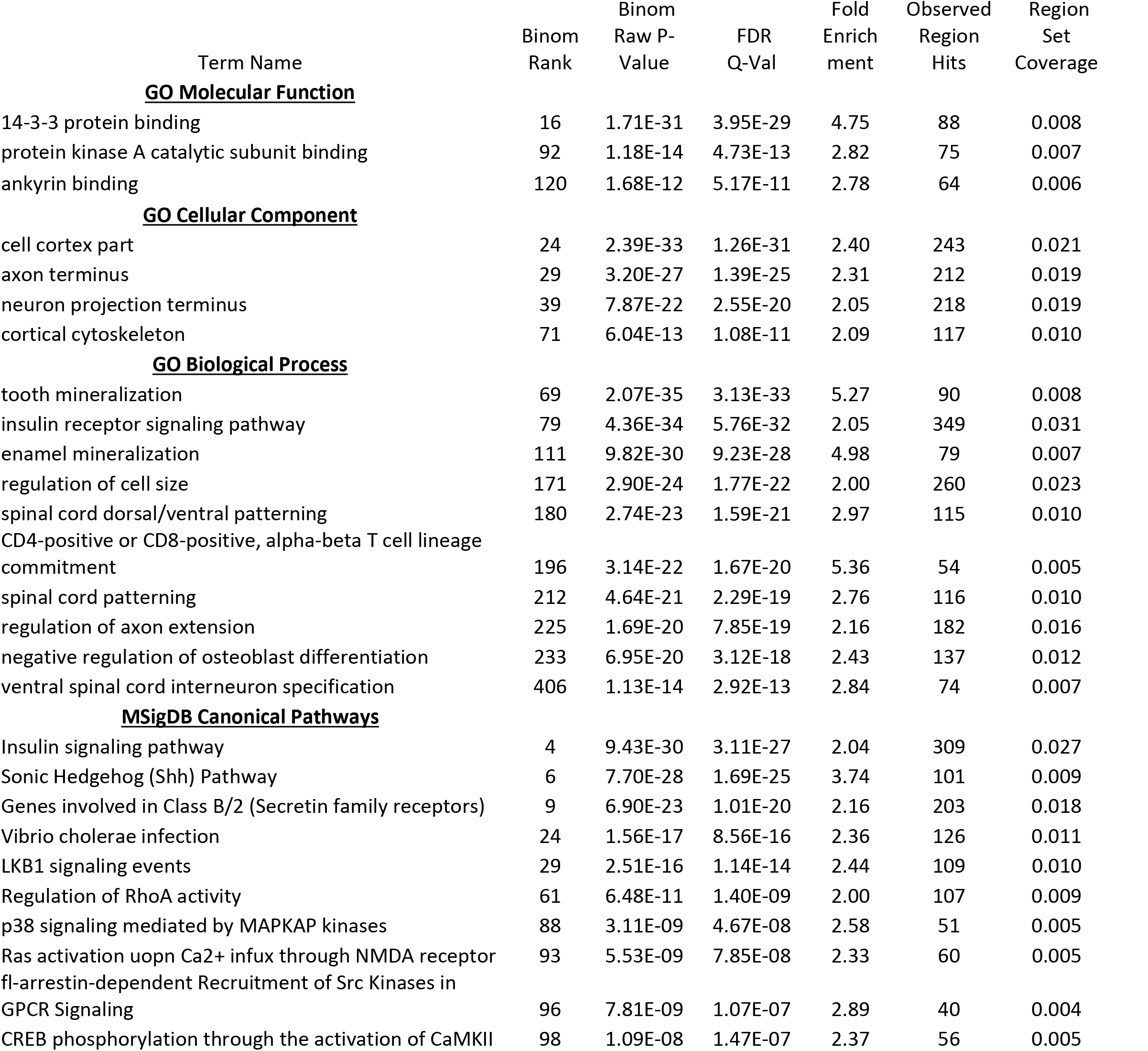
Functional annotation and pathway analysis of TCF4 genomic binding sites using GREAT.

The full set of 11,322 TCF4 binding sites implicated 6528 unique genes ±10 kb of the gene body (Supplementary Table S2), which is approximately one quarter of all genes in refGene, including non-coding RNAs and genes with provisional nomenclature designations. We used FUMA (27) to test for enrichment in tissue-specific differentially expressed gene sets (Supplementary Figure S7). The TCF4 gene set was most enriched in genes up-regulated in the brain and pituitary, and genes down-regulated in the heart and blood vessels. To further probe the expression patterns of the TCF4 gene set, we looked at single cell RNA-seq data for specific CNS cell types. We observed that the TCF4 gene set was significantly over-expressed in pyramidal neurons from the somatosensory cortex (p = 5.2x10^−5^, Figure 3).

**Figure 3.**
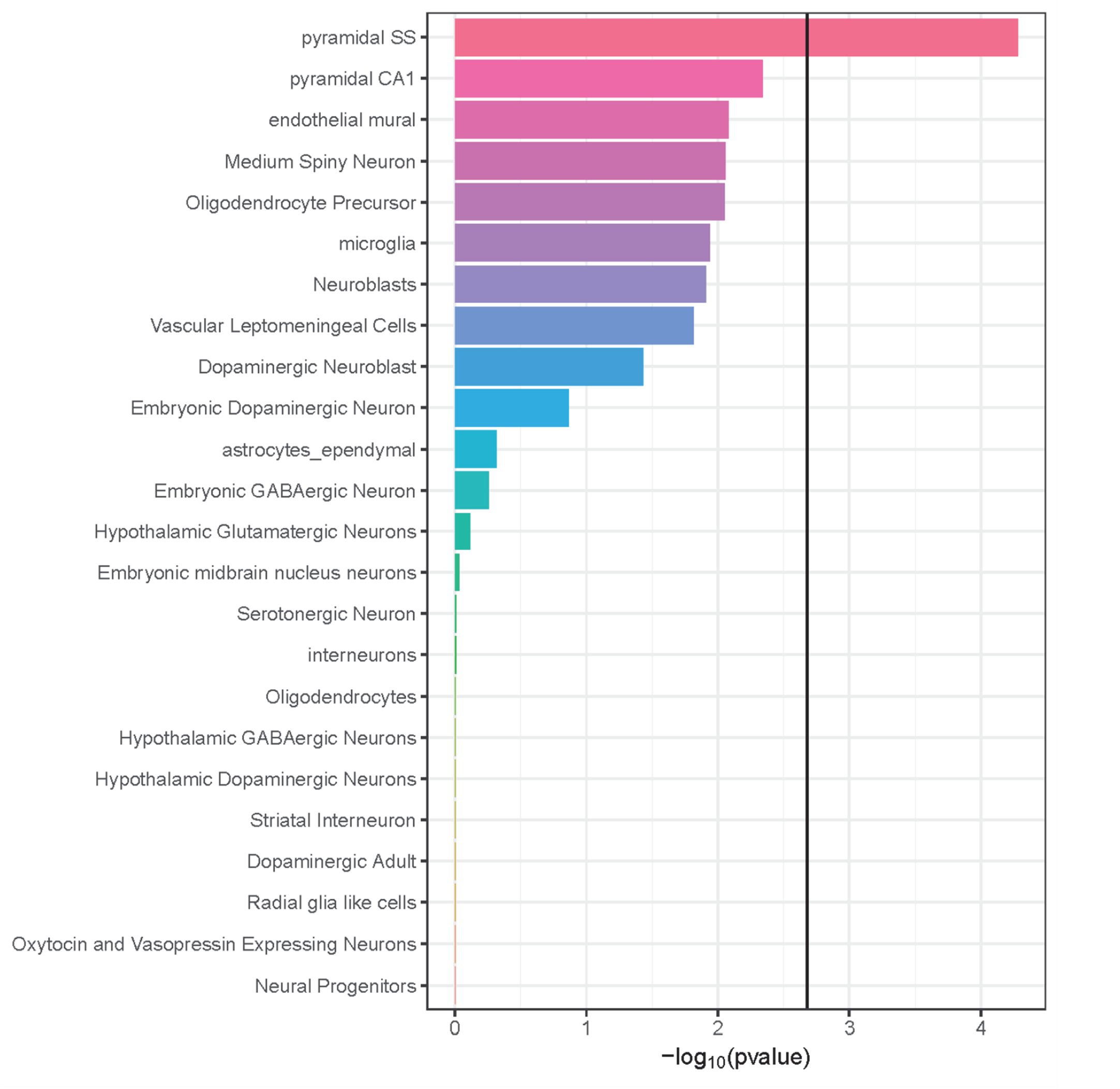
Enrichment of TCF4 gene set expression in specific brain cell types as determined by single cell RNA-seq (47). We tested if expression of genes in the TCF4 gene set were significantly higher than for genes not in the gene set for each cell type. The bold line at ∼2.7 on the x-axis is the Bonferroni-adjusted –log_10_(p-value) for multiple testing (α=0.05/24). “Pyramidal SS” = pyramidal neurons of the somatosensory cortex. “Pyramidal CA1” = pyramidal neurons of the CA1 region of the hippocampus.

### Differential regulation of TCF4 genes in siRNA knockdown experiments and post-mortem data

We integrated our ChIP-seq data with relevant datasets to refine our findings and identify SZ risk genes under TCF4 control. We first tested for overlap between the TCF4 gene set and gene expression data from a *TCF4* knockdown experiment using small interfering RNA (siRNA), also conducted in SH-SY5Y cells (28). We expected that genes differentially expressed following *TCF4* knockdown should be enriched for those with TCF4 binding sites from ChIP-seq. A significant enrichment was observed and this was driven by genes downregulated following TCF4 knockdown (Table 2). No significant enrichment was observed in the upregulated genes. We also tested the TCF4 gene set for enrichment in a second TCF4 siRNA knockdown experiment, this time in cortical neuron progenitor cells (29) (Table 2). Once again downregulated genes, but not upregulated genes, were significantly enriched. This consistency was encouraging. All genes overlapping between ChIP-seq and siRNA studies are provided in Supplementary Tables S4 and S5.

**Table 2.**
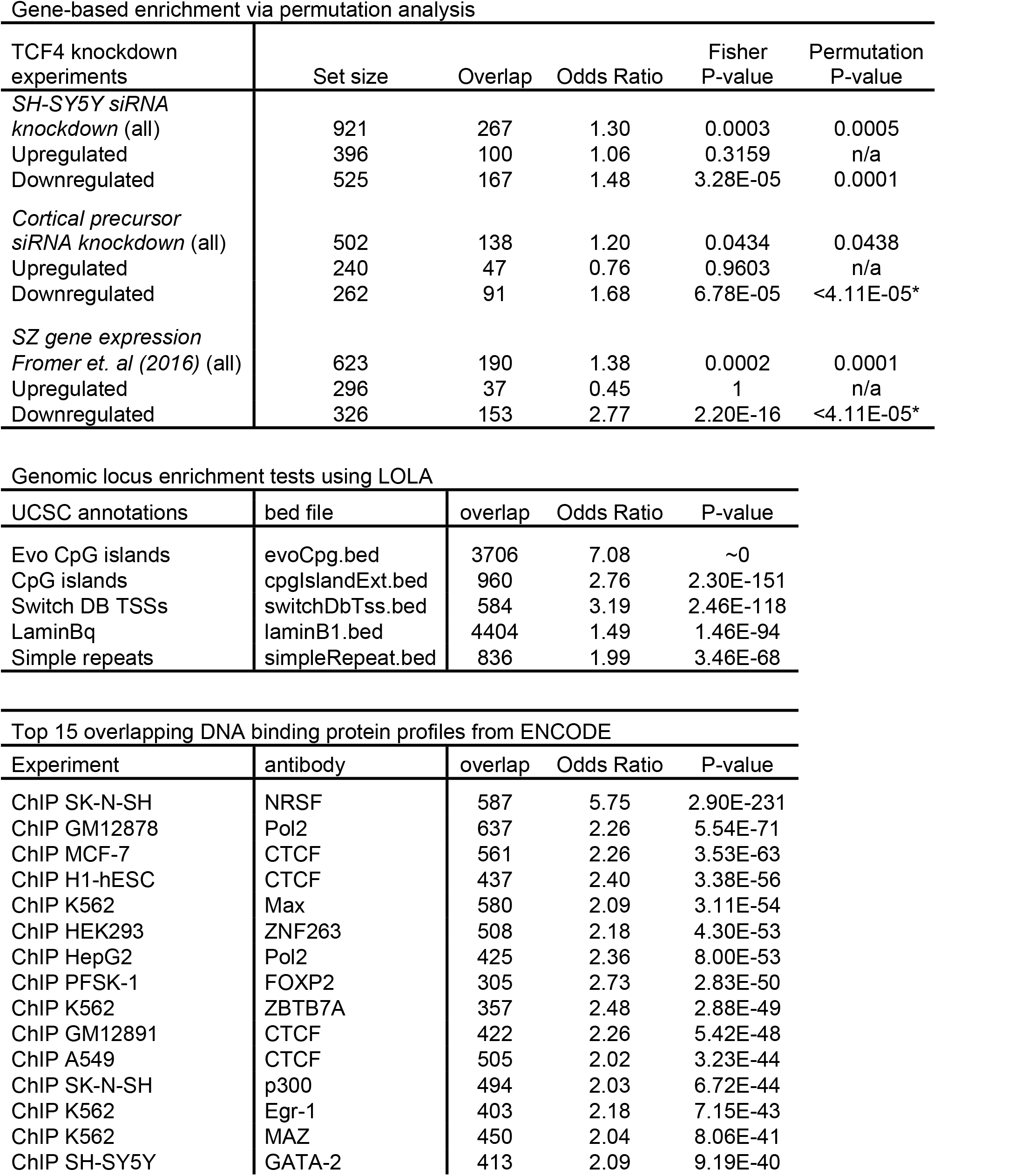
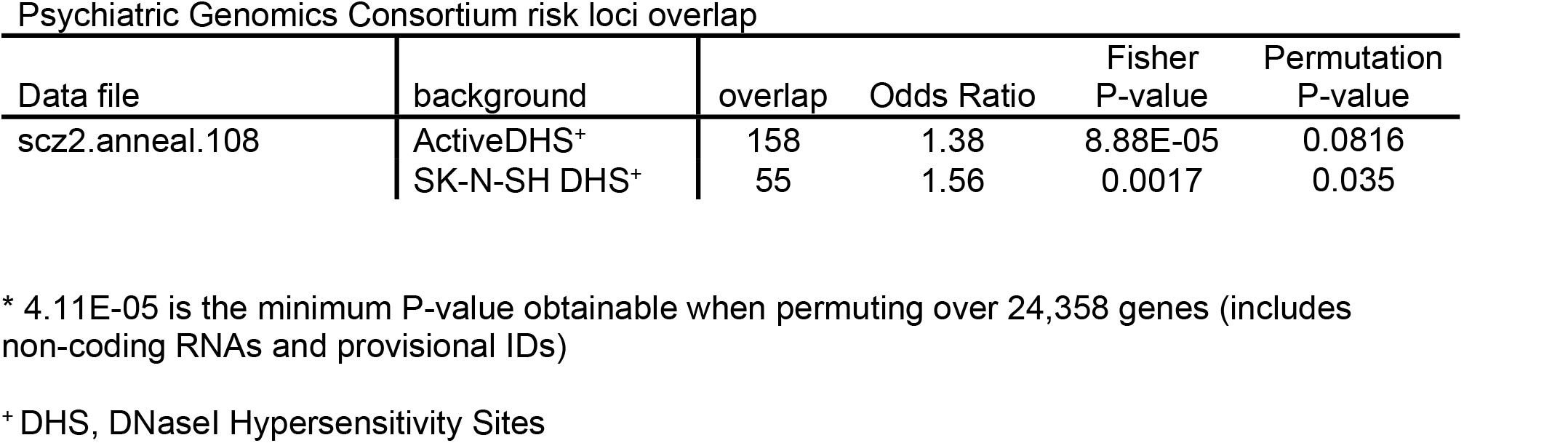
Integration of TCF4 binding profile with functional and SZ datasets.

We next tested if the TCF4 gene set was enriched among genes differentially expressed in post-mortem brain tissue from SZ patients. Here we used findings from Fromer et al. (2016) (30), who conducted bulk RNA-seq of dorsolateral prefrontal cortex (DLPFC) from 258 SZ cases and 279 controls. Once again, downregulated genes were significantly enriched with almost half possessing a TCF4 binding site, compared to just 12.5% of the up-regulated genes. This highly significant enrichment in down-regulated genes was surprising because these are downregulated in SZ, not as a direct result of *TCF4* knockdown. We hypothesized that *TCF4* is driving the down-regulation of genes in SZ by being downregulated itself, analogous to the situation in the siRNA experiments. However, *TCF4* expression was significantly *upregulated* in the SZ patients (fold enrichment = 1.16, Fromer et. al, their Supplementary Data File 3 (30)). Further analysis of these gene sets showed that few genes were shared in common. That is, only seven genes with TCF4 binding sites from ChIP-seq were downregulated in the SH-SY5Y siRNA study (28) *<u>and</u>* down-regulated in the SZ expression study (30). These were *ANKMY1, FAM78A, IGF2, MXRA8, NT5M, PELI3* and *TNS3*. Notably, *IGF2* had the largest fold-change reduction of all genes in the Fromer et al. study. A further two genes with TCF4 binding sites, *DBNL* and *PNPLA7*, showed downregulation in the neural progenitor cell siRNA study (29) *and* the SZ gene expression study.

### Overlap of TCF4 binding sites with other TFs and SZ risk loci

We next considered overlap of TCF4 binding sites with UCSC genome browser features. TCF4 binding sites were highly enriched (OR=7.08) in CpG islands, classified according to the Weizmann Institute CpG evolution model (31), and in transcription start sites (OR=3.19) from the SwitchGear Genomics library (www.switchgeargenomics.com) (Table 2). We also looked at overlap with binding profiles for other TFs from ENCODE. Notable overlapping factors were FOXP2, a neural transcription factor involved in speech development(32), and p300, a chromatin remodeler encoded by the *EP300* gene that is associated with SZ (1).

Our final data integration analysis was with the 108 PGC2 SZ risk loci published in 2014 (1). For this analysis, we used genomic locus-based enrichment rather than gene sets. This is important because many SZ-associated genes are large and may be more likely to overlap genomic annotations by chance than a randomly selected gene set (33). Therefore, annotations can be confounded with gene size, leading to erroneous conclusions of enrichment. Testing for TCF4 binding site enrichment in the associated genomic regions, and not considering occurrence in genes, obviates this potential bias. Overall, 39 of the 108 PGC SZ loci contained one or more TCF4 binding sites, with 130 sites in total falling within their boundaries. Permutation testing of enrichment using all known human regulatory regions as the background set revealed a non-significant overlap (p=0.082). Narrowing the background set to relevant regulatory regions for the cell type used in the ChIP-seq experiments revealed a nominally significant enrichment (p=0.035) (Table 1). PGC SZ genes with TCF4 binding sites ±10 kb that were also differentially expressed in either of the siRNA experiments described above were *APH1A, C1orf54, CENPM, CHRNA5, DFNA5, GFOD2, GRAMD1B, LRP1, MPP6, PDCD11, SEZ6L2, TLE1*, and *XRCC3*. Of these, both *LRP1* and *XRCC3* had three binding sites in our ChIP-seq data, *PDCD11* had two, while the remainder had one each.

### The TCF4 isoform recognized by our antibodies is down-regulated in differentiated neurons

Pathways and specific genes genes among our findings suggested that TCF4-regulated genes in our study are involved in neuronal growth and/or differentiation. To determine if there are developmental differences in expression of the TCF4 isoform we captured in ChIP-seq, we differentiated LUHMES neural precursor cells using an established protocol that yields post-mitotic neurons in 5 days (34) (Figure 4). We used Western blotting with the N-16 antibody to probe for TCF4 in cell lysates and compared SH-SY5Y, undifferentiated LUHMES and differentiated LUHMES cells. Figure 4 shows a reduction in the TCF4 signal in the differentiated cells. Quantitative analysis of three duplicate experiments indicated that the TCF4 isoform recognized by N-16 was expressed in the differentiated cells at <10% of the levels in the undifferentiated, rapidly growing cells.

**Figure 4.**
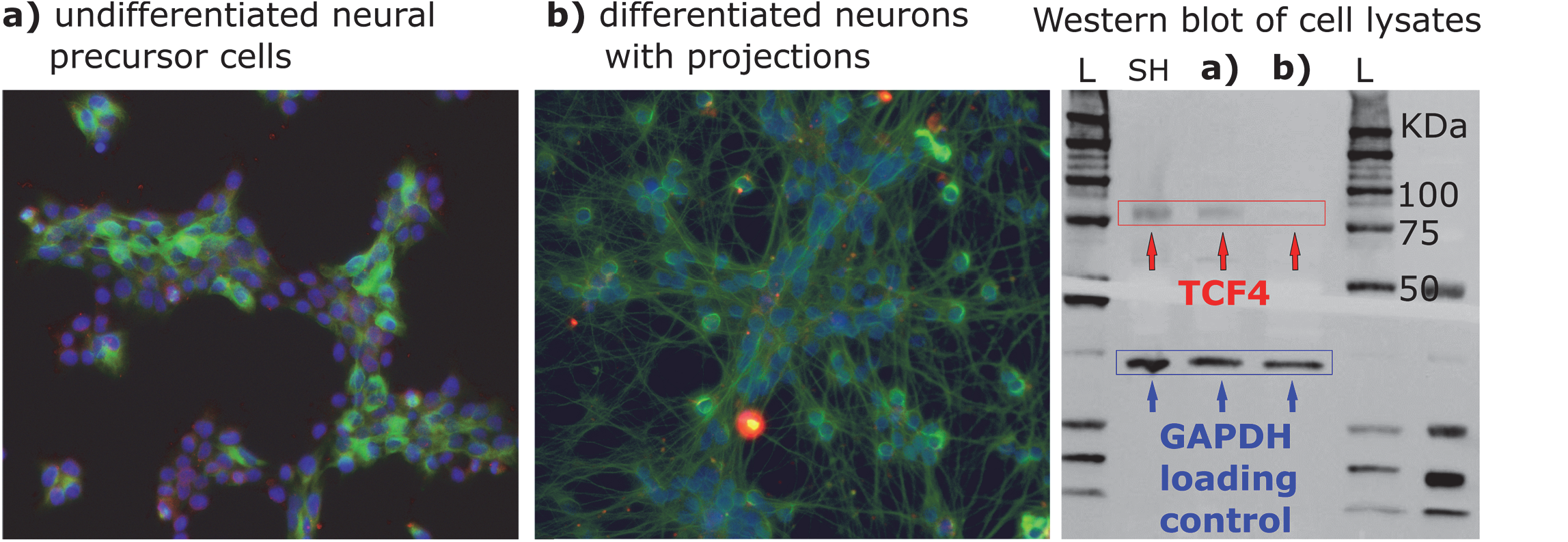
Differentiation of LUHMES neuronal precursor cells (a) to mature neurons (b) in culture. The third panel shows Western blotting for TCF4 using lysates from SH-SY5Y (SH), undifferentiated (a) and differentiated (b) LUHMES. DAPI stain (blue) was used to identify cell bodies, while neuronal projections were stained using anti-beta tubulin antibody (green). The Western image (panel c) shows that TCF4 is down-regulated in the differentiated LUHMES neurons.

## DISCUSSION

We obtained data on TCF4 binding from ChIP-seq, which allowed us to probe the TCF4 gene network for association with SZ risk genes. We validated our antibodies according to ENCODE standards. Following ChIP-seq, we further validated our findings by showing that our TCF4 binding sites were enriched for the known TCF4 Ebox binding motif. We also showed that genes with TCF4 binding sites were enriched among genes differentially expressed in TCF4 siRNA knockdown experiments. Taken together, these results strongly support the validity of the data.

A limitation of our ChIP-seq experiments was that only two out of three antibodies yielded usable data. It is accepted that antibodies that pass initial characterization may still fail to yield good ChIP-seq data (25), yet the immunoblot for the antibody that failed (1G4) is arguably the cleanest (Figure 1b). However, antibody 1G4 is monoclonal as compared to K-12 and N-16 that are polyclonal. While monoclonal antibodies have advantages in specificity and reproducibility, they are more prone to fail in ChIP-seq because the single epitope in the protein may be obscured by the bound DNA. Another limitation is that we only have data on the long TCF4 isoform (TCF4-B). The *TCF4* locus can give rise to several distinct isoforms via alternative splicing (18). Some isoforms are exclusively nuclear while others rely on heterodimerization partners. Understanding the roles of these variants in the CNS will require additional research.

Several genes that we identified as having TCF4 binding sites have been functionally linked with TCF4 in prior studies. For example, TCF4 has also been shown to affect neural excitability via repression of potassium channel *KCNQ1* (11). In our data, this gene had 19 unique TCF4 binding sites, the 16^th^ largest number for any gene. Conversely, given the problems with “TCF4” nomenclature, delineating what is *not* seen in our data may be of value. Several articles describe the interaction between “TCF4”, β-catenin and p300 (35). This relates to T Cell Factor 4, encoded by *TCF7L2*. Betacatenin is encoded by *CTNNB1* and we did not detect a TCF4 binding site at this gene. Furthermore, even though the binding profiles of p300 (encoded by *EP300)* and TCF4 strongly overlap (Table 2), TCF4 does not appear to regulate *EP300*, at least in our data. The co-occurrence of their binding sites does, however, imply that they may be involved in the regulation of a partially overlapping set of genes in CNS cells. *EP300* is a PGC2 SZ risk locus (1) and was associated with emotional processing in functional neuroimaging experiments (36). Further analysis into the overlapping pathways regulated by these two SZ-associated DNA binding proteins may be relevant.

Several genes with TCF4 binding sites in our ChIP-seq data were down-regulated in *TCF4* siRNA knockdown experiments. A subset of these genes were also dysregulated in post-mortem brain tissue from SZ patients. Among these, *IGF2* (insulin-like growth factor 2) was the most down-regulated gene in postmortem SZ brain in the study by Fromer *et al*. (30). *IGF2* may regulate neural plasticity to modulate behavior and memory (37). Furthermore, deficits in hippocampal neurogenesis in a mouse model of 22q11.2 deletion-associated SZ can be rescued by *IGF2* (38). A significant amount of work has been conducted on the role of *IGF2* in the brain, yet few studies address the role of *IGF2* in SZ etiology. One issue is that *IGF2* was not detected among PGC SZ risk loci (1). This may indicate that down-regulation of *IGF2* in SZ is a consequence of risk variants at other loci or environmental factors, rather than as a result of risk variants at the *IGF2* locus itself.

Several PGC SZ risk genes contained TCF4 binding sites and thirteen of these also showed differential expression in *TCF4* siRNA experiments. The risk loci identified by the PGC may span several hundred kb and contain many genes. It is often not apparent which genes in these regions should be selected for further study. For example, there are 14 unique genes in the third most significant region identified by the PGC (chr10:104423800-105165583, P-value = 6.12E-19) (1). Among these, *PDCD11* is likely regulated by TCF4, exhibiting both TCF4 binding sites and differential expression in a *TCF4* siRNA study. It is also expressed in brain (39) but is poorly characterized. Its likely regulation by TCF4 may suggest further work is merited to characterize this gene.

Many of the PGC SZ risk genes regulated by TCF4 are involved in neuronal differentiation and development. For example, *APH1A* encodes a subunit of gamma secretase, is crucial for Notch signaling in embryogenesis and is predominantly expressed in non-neuronal and neuronal precursor cells (40). *LRP1* has diverse roles in the CNS and is a regulator of neural progenitor cell function (41), while *TLE1* functions as a transcriptional repressor to regulate neuronal differentiation (42). Furthermore, we observed that the specific TCF4 isoform captured in our ChIP-seq experiments is up-regulated in rapidly growing, undifferentiated neural precursors relative to differentiated neurons. We conclude that we have identified potential regulatory interactions between a SZ-associated TF and several SZ risk loci that implicate processes involved in neuronal development. As an “omic” study, our findings represent hypotheses to be tested in future neurobiological studies. We hope that the mapping of these interactions will stimulate new research into TCF4.

## MATERIALS AND METHODS

### Antibody selection

At the start of this study, no ChIP-grade antibodies were available for TCF4, so we selected candidate antibodies from commercial vendors. Due to the confusion in nomenclature with T-Cell Factor 4, we discovered several mislabeled antibodies. Therefore, we limited ourselves to antibodies with published epitope sequences that could be confirmed as TCF4 via protein BLAST (NCBI). Eight antibodies were selected, of which three passed initial QC and were used for ChIP-seq: polyclonal anti-TCF4 antibodies K-12 (sc-48947) and N-16 (sc-48949) from Santa Cruz Biotechnology (Dallas, TX) and monoclonal anti-TCF4 antibody 1G4 (Novus, Littleton, CO).

### Cell culture

SH-SY5Y cells (ATCC, Manassas, VA) were cultured according to supplier’s standard protocols. Cell line authentication via simple tandem repeat genotyping was conducted by the University of Arizona Genomics Core Facility. LUHMES cells were cultured and differentiated according to a published protocol (34). Additional details and immunocytochemistry methods are in the Supplementary Methods.

### Antibody validation

We followed ENCODE guidelines for antibody validation (25). In addition to Western blotting of SH-SY5Y cell lysates, we used immunoprecipitation (IP) of TCF4 protein followed by Western blotting or protein mass spectrometry to characterize the proteins captured in IP (see Supplementary Methods). Further validation after ChIP-seq used motif enrichment testing of binding site sequences (500 bp, centered on ChIP-seq peaks) with using DREME with default settings as implemented in MEME-ChIP (43).

### Chromatin Immunoprecipitation sequencing (ChIP-seq)

Each ChIP assay used approximately 1.2x10^7^ SH-SY5Y cells and was performed with the SOLiD ChIP-Seq Kit (Life Technologies) according to manufacturer’s specifications, with some adjustments (Supplementary Methods). ChIP-Seq libraries were validated using the BioAnalyzer high-sensitivity chip assay (Agilent) prior to multiplexed high throughput sequencing on the SOLiD 5500 platform. 50 bp single end reads were generated, with a target read number of 25 million tags per sample. In addition to ChIP samples using anti-TCF4 antibodies, their respective IgG controls and input DNA controls were sequenced.

### ChIP-seq data analysis

Reads were aligned to the human genome (build hg19) using BioScope 1.2 (Life Technologies). Multi-mapping reads were discarded and only stringent single alignments retained. Sample files were output as .bam files using BioScope. PCR duplicates were removed by dropping multiple reads with identical start positions using the Samtools (44) rmdup function and alignment files were written in tagAlign.gz format using Samtools and Bedtools (45). To call peaks and assess experiment quality, we used SPP (21) distributed with phantompeakqualtools (46). A false discovery rate (FDR) threshold of 1%, as implemented in SPP, was used to call peaks. Any peaks that mapped to hg19 ENCODE blacklisted regions (ftp://encodeftp.cse.ucsc.edu/users/akundaje/rawdata/blacklists/hg19/wgEncodeHg19ConsensusSignalArtifactRegions.bed.gz) were removed.

### Data Integration

Bioinformatics analysis was conducted in R (www.r-project.org). Gene lists were obtained from refGene via UCSC Genome Browser download (August 10^th^ 2017), followed by elimination of transcripts with ambiguous mapping and pruning entries by maximum boundary to yield a single non-redundant locus per gene. Gene pathway analysis used GREAT(26) version 3.0.0, assigning proximal regulatory domains ± 10 kb. Tissue-specific expression of the top TCF4 genes was evaluated using the “Gene2Func” mode in FUMA (27).

For cell-specific expression analysis, we obtained single cell RNA-seq data from five brain regions in mice (9970 single cells) that were previously clustered into 24 different cell types (47). Normalization factors were computed for each of the 9970 single cells using the scran R package (48, 49) using the 50% of the genes with mean expression higher than the median. The normalization factors were computed after clustering cells using the scran quickcluster() function to account for cell type heterogeneity. We then performed 24 differential expression analyses using BPSC (50) testing each cell type against the 23 other cell types using the normalization factor as a covariate. For each differential expression analysis, the t-statistics were then transformed to a standard normal distribution. Finally, for each cell type, we used linear regression to test if the standard normalized t-statistics for genes in the TCF4 gene set were significantly higher than for genes not in the gene set.

Enrichment testing of TCF4 binding sites in significant genes from siRNA knockdown expression studies (28, 29) was carried out by mapping genes ± 10 kb, followed by one-sided Fisher exact tests. Permutation testing of significant findings used the shiftR package (https://github.com/andreyshabalin/shiftR), as outlined previously (51). Test of overlap between TCF4 peaks and genomic annotations used the LOLA Bioconductor package (52), with all mappings obtained from the LOLACore annotation set (databio.org/regiondb). The background set for this analysis was the default DNase hypersensitive sites (DHS) in multiple tissues (“activeDHS” set), which captures known human regulatory regions (52, 53). Testing for overlap with PGC SZ findings was also based on genomic locus, rather than gene. SZ-associated loci were obtained by download of the “scz2.anneal.108” file from the PGC (https://www.med.unc.edu/pgc/results-and-downloads). Background sets for this analysis were either “activeDHS” as above or DNase hypersensitive sites specifically for SK-N-SH cells (“wgEncodeOpenChromDnaseSknshPk” track, Duke DHS from ENCODE). No DHS data for SH-SY5Y were available, but SH-SY5Y are a subline of SK-N-SH isolated from the same donor (54). After matching to background set, enrichment testing used Fisher exact tests followed by permutation as above.

## Acknowledgements

Cell Line Authentication was provided by Elizabeth Cox and the team at University of Arizona Genetics Core (via Science Exchange). Thanks to Derek Blake at the MRC Centre for Neuropsychiatric Genetics and Genomics, Cardiff University, United Kingdom for providing siRNA gene expression data, and to Jens Hjerling-Leffler, Sten Linnarsson, Ana Munoz Manchado and Amit Zeisel of the Karolinska Institute, Stockholm, Sweden for providing single-cell RNA-sequencing data. This article was submitted as a preprint to bioRxiv (11/07/2017) and assigned doi: https://doi.org/10.1101/215715. This work was funded through grant R21 MH099419 from the US National Institute of Mental Health to JL McClay.

## Conflict of Interest statement

The authors declare no financial conflicts of interest.

## Web resources

All raw data (tagAlign.gz files) and accompanying descriptions are available at the Gene Expression Omnibus (GEO) repository. *(Submitted at time of submission and accession number pending)*

